# MpILR1 Hydrolyzes Jasmonate-Amino Acid Conjugates to Activate dn-*iso*-OPDA Signaling in *Marchantia polymorpha*

**DOI:** 10.64898/2026.09.24.754050

**Authors:** Wenting Liang, Ángel M. Zamarreño, Takuya Kaji, Runa Tanaka, Harushi Suzuki, Minoru Ueda, José M. García-Mina, Roberto Solano, Andrea Chini

## Abstract

Jasmonates are essential phytohormones that coordinate defense responses and developmental programs across land plants. In angiosperms, the active jasmonate ligand jasmonoyl-L-isoleucine (JA-Ile), is produced through GH3-mediated conjugation of jasmonic acid to isoleucine and JA-Ile homeostasis is further shaped by ILR1/ILL-family amidohydrolases. In contrast, the primary bioactive jasmonate ligand in bryophytes, dinor-12-oxo-phytodienoic acid (dn-*iso*-OPDA), is inactivated through conjugation with amino acids, raising the question of whether these conjugates constitute a reversible hormone reservoir or an irreversible catabolic end point. Although the ILR1-like family has been characterized extensively for its role in auxin and jasmonate homeostasis in angiosperms, its function in bryophytes remains basically unexplored. Here we show that MpILR1, the sole Marchantia ortholog of the ILR1/ILL family, hydrolyzes a specific subset of dn-*iso*-OPDA-amino acid conjugates *in vivo*. Loss-of-function Mp*ilr1* mutants exhibit enhanced accumulation of dn-*iso*-OPDA conjugated to hydrophobic amino acids (Val, Leu and Ile) but not to hydrophilic residues (His, Glu and Gln), demonstrating substrate-selective hydrolysis. MpILR1 hydrolytic activity is required for full dn-*iso*-OPDA-mediated responses, including transcriptional activation and defense against gastropod herbivory. These findings establish MpILR1 as a key positive regulator of jasmonate signaling in *Marchantia polymorpha* and suggest that hormone conjugation/deconjugation is an ancient regulatory mechanism evolved during plant terrestrialization.

## Introduction

Jasmonates (JAs) are essential phytohormones that coordinate plant defense against mechanical wounding, herbivory, and necrotrophic pathogen infection, while also regulating growth and developmental transitions (Howe et al., 2018; Gasperini and Howe, 2024). Land plants rely on a conserved jasmonate signaling module to translate mechanical and biotic stress into transcriptional defense output; the evolutionary history of this pathway offers a unique opportunity to infer how hormone specificity and homeostatic regulation co-evolved across land plant diversification (Bowman et al., 2017; Howe et al., 2018; Monte et al., 2018). In flowering plants, jasmonic acid (JA) is mainly synthesized from α-linolenic acid through the well-characterized octadecanoid pathway, in which 12-oxo-phytodienoic acid (OPDA) serves as a precursor of JA, which is subsequently conjugated to L-isoleucine by GRETCHEN HAGEN 3 (GH3) enzymes to form the bioactive hormone jasmonoyl-L-isoleucine (JA-Ile) (Staswick and Tiryaki, 2004; Fonseca et al., 2009; Delfin et al., 2022). JA-Ile perception by the Coronatine insensitive 1 (COI1)-Jasmonate ZIM-domain (JAZ) co-receptor complex initiates ubiquitin-mediated proteolysis of JAZ repressors, releasing transcription factors that activate genome-wide defense responses (Chini et al., 2007; Thines et al., 2007; Sheard et al., 2010; Chini et al., 2016).

A recurring theme in phytohormone biology is the involvement of amino acid conjugation and hydrolysis in the regulation of phytohormones, suggesting that it may represents a broadly conserved mechanism for controlling hormone availability and to fine-tune signaling amplitude (Jez, 2022). For auxin, GH3-catalyzed conjugation of indole-3-acetic acid (IAA) generally produces storage inactive IAA amino acid conjugates; in contrast, the reverse reaction, hydrolysis mediated by amidohydrolases of the ILR1 (IAA-Leucine Resistant 1)-LIKE (ILL) family, releases bioactive IAA and consequently activate auxin signaling (Bartel and Fink, 1995; LeClere et al., 2002). A similar GH3/ILR-mediated conjugation-hydrolysis system also regulates *cis*-OPDA/dn-*cis-*OPDA, a JA-Ile precursor that also activates signaling output independently of its conversion to JA-Ile (Chini et al., 2018; Monte et al., 2020). In Arabidopsis, *cis*-OPDA is conjugated to several amino acids generating conjugates that predominantly function as temporary storage, as their hydrolysis by ILR1/ILL-family amidohydrolases can restore the availability of free *cis*-OPDA during stress responses (Široká et al., 2025). The homeostasis of the bioactive jasmonate JA-Ile is also regulated by conjugation-hydrolysis cycle, but with the opposite functional polarity. Conjugation of JA to isoleucine and additional amino acids is the activating step, whereas hydrolysis of JA-Ile by ILL-family amidohydrolases, principally IAR3 (Jasmonate Responsive 3) and ILL6 in Arabidopsis, decreases the levels of bioactive JA-Ile and thereby attenuates the signal pathway (Widemann et al., 2013; Woldemariam et al., 2012; Zhang et al., 2016). The same core enzymatic module, GH3-type conjugation paired with ILR1/ILL-type deconjugation, has therefore been conserved through evolution to fine-tune different regulatory pathways, underscoring the mechanistic flexibility of amino acid conjugation as a general strategy for hormone homeostasis in plants (Westfall et al., 2012; Stepanova and Alonso, 2016).

This dual regulatory logic raises an important evolutionary question on whether the conjugation-hydrolysis process was already present and already coupled to defense hormone regulation in the first land plant lineages. The liverwort model plant *Marchantia polymorpha* offers a unique opportunity to address this question, because it lacks JA-Ile, the canonical bioactive ligand in euphyllophyte plants (major vascular plant clade encompassing ferns, horsetails and angiosperms), and instead relies on dn-*iso*-OPDA as the endogenous ligand of a functionally conserved MpCOI1-MpJAZ co-receptor complex (Monte et al., 2018; Monte et al., 2019; Peñuelas et al., 2019; Chini et al., 2023). Our previous work established that, contrary to jasmonates in Arabidopsis, MpGH3A conjugates bioactive dn-*iso*-OPDA to a range of amino acids, generating derivatives with reduced signaling activity, most abundantly dn-OPDA-His, -Glu and -Gln (Liang et al., 2025). These results demonstrate that a conjugation-based inactivation mechanism regulates the bioactive dn-*iso*-OPDA in Marchantia, paralleling the auxin-inactivating role of GH3 conjugation in angiosperms.

However, whether Marchantia also possesses the complementary hydrolytic branch of this cycle, and whether such hydrolysis would activate dn-*iso*-OPDA signal, has not been explored. Addressing this question will clarify whether dn-*iso*-OPDA-amino acid conjugates constitute a genuine, dynamically regulated hormone reservoir or instead represent catabolic end products with no further regulatory function. In addition, Marchantia encodes only a single *ILR1/ILL* ortholog, in contrast to the functionally diversified seven-member family in Arabidopsis (Bowman et al., 2017); characterizing this single hydrolase offers an excellent opportunity to infer the ancestral physiological function of this hydrolase family, prior to its expansion and functional diversification in vascular plants. Here, we address this gap by generating loss-of-function mutants of MpILR1 and studying its role in dn-*iso*-OPDA conjugate turnover, hormone homeostasis and defense responses in *Marchantia polymorpha*. We observed that hydrolysis of specific dn-*iso*-OPDA conjugates is a naturally occurring process and that MpILR1 is required for the hydrolysis of a specific subset of dn-*iso*-OPDA conjugates. Furthermore, we showed that MpILR1-mediated conjugate turnover contributes to the accumulation of bioactive dn-*iso*-OPDA and to full activation of MpCOI1-dependent anti-herbivore defenses.

## Materials and methods

### Plant material

The *MpILR1* gene was targeted via CRISPR-Cas9 nickase-mediated mutagenesis using the *Marchantia polymorpha* Tak-1 accession as the wild-type background. To target the first exon of the gene, four pairs of gRNAs (eight in total; Supplementary Table S1) were designed and initially cloned into the vectors pBC-GE12, pBC-GE23, pBC-GE34, and pMPGE_EN04. These gRNAs were subsequently transferred via an LR reaction into the pMpGE017 binary vector, which expresses the CRISPR-Cas9 nickase. Transformation was performed on WT plants using the regenerating thalli method, and successful transformants were selected based on hygromycin resistance (Kubota et al. 2013). Candidate mutants were selected by extracting and sequencing genomic DNA from hygromycin-resistant explants as previously described (Soriano et al., 2022; Liang et al., 2025). In addition, the Mp*coi1-2* mutant plants were used as control in experiments analyzing OPDA-mediated responses (Kaji et al., 2024; Liang et al., 2025).

### Culture conditions and wounding

Plants were routinely cultured on half-strength Gamborg’s B5 medium supplemented with 1% agar at 21 °C under continuous white light (50–60 µmol m^−2^ s^−1^). For quantification of jasmonate-derived molecules, thalli were grown for 2 weeks prior to analysis. Mechanical wounding was performed on both wild-type and mutant thalli by crushing the entire plants with tweezers, following established protocols (Kneeshaw et al. 2022; Soriano et al. 2022). At the indicated time points, the wounded tissues were harvested and immediately flash-frozen in liquid nitrogen.

### Growth inhibition assays

Growth inhibition assays were conducted following established procedures (Kneeshaw et al., 2022; Soriano et al., 2022; Liang et al., 2025). Briefly, half-strength Gamborg’s B5 (GB5) medium was supplemented with 10 μM OPDA-amino acids. After approximately 2 weeks of growth, plant images were captured using a Nikon D1-x digital camera, and the surface area of each plant was quantified using ImageJ software. Relative growth performance was determined by calculating the ratio of the surface area of hormone-treated plants to that of the untreated controls. All experiments were performed at least three times with consistent results.

### Chemicals

All chemical reagents and solvents were obtained from commercial suppliers (Kanto Chemical Co. Ltd., Wako Pure Chemical Industries Co. Ltd., Nacalai Tesque Co. Ltd., Tokyo Chemical Industry Co. Ltd., Sigma-Aldrich Co. LLC.) and used without further purification. All anhydrous solvents were either dried by standard techniques and freshly distilled before use or purchased in anhydrous form and used as supplied. Reversed-phase high–performance liquid chromatography (HPLC) was carried out on a PU–4180 plus pump equipped with UV–4075 and MD–4010 detectors (JASCO, Tokyo, Japan). 1H and 13C NMR spectra were recorded on a JNM–ECS–400 spectrometer (JEOL, Tokyo, Japan) in deuterated chloroform using TMS as an internal standard. Fourier transform infrared (FT/IR) spectra were recorded on an FT/IR–4100 (JASCO, Tokyo, Japan). High–resolution (HR) electrospray ionization (ESI)–mass spectrometry (MS) analyses were conducted using a microTOF II (Bruker Daltonics Inc., MA, USA). Optical rotations were measured using a JASCO P–2200 polarimeter (JASCO, Tokyo, Japan). Flash chromatography was performed on an Isolera system (Biotage Ltd., North Carolina, USA). TLC analyses were performed on Silica gel F254 (0.25 mm or 0.5 mm, MERCK, Germany). All reactions were carried out under air unless stated otherwise.

### Synthesis of dn-*iso*-OPDA-Leu

To a solution of dn-*iso*-OPDA (49.1 mg, 0.19 mmol) and Et_3_N (80.0 µL, 0.58 mmol) in THF (2.0 mL) was added ethyl chloroformate (38.0 µL, 0.40 mmol) at 0 °C under an argon atmosphere. After stirring at room temperature for 2 h, the above mixture was transferred to a solution of L-leucine (127 mg, 0.97 mmol) and diisopropylethylamine (200 µL, 1.17 mmol) in H_2_O (2.0 mL) and the resulting mixture was stirred for 3 h at room temperature. The reaction mixture was then acidified with 1 M aq. HCl and extracted with CHCl_3_. The combined organic layer was washed with brine, dried over Na_2_SO_4_, and concentrated under reduced pressure. The residue was purified by silica gel medium-pressure chromatography (CHCl_3_/MeOH/AcOH = 99/1/0.1 to CHCl_3_/MeOH/AcOH = 90/10/0.1) to give dn-*iso*-OPDA-Leu (63.3 mg, 89%) as a pale yellow oil. The obtained dn-*iso*-OPDA-Leu (24.9 mg) (Supplemental Figure S1 and S2) was further purified by RP-HPLC (ODS-HG-5 column: F 20 × 250 mm, Flow rate 8.0 mL/min, detection 220 nm, eluent: MeOH/H_2_O/AcOH = 80/20/0.1, t_R_ = 11.4 min) to give dn-*iso*-OPDA-Leu (11.1 mg) as a colorless oil.

[α]_D_^25^ = −0.72 (c 0.54, CHCl_3_). ^1^H-NMR (400 MHz, CDCl_3_) δ_H_: 5.96 (d, *J* = 7.8 Hz, 1H), 5.36 (dtt, *J* = 10.7, 7.3, 1.7 Hz, 1H), 5.19 (dtt, *J* = 10.7, 7.1, 1.6 Hz, 1H), 4.59 (m, 1H), 2.92 (d, *J* = 7.1 Hz, 2H), 2.49-2.48 (m, 2H), 2.43 (t, *J* = 7.6 Hz, 2H), 2.38-2.36 (m, 2H), 2.25 (td, *J* = 7.4, 4.7 Hz, 2H), 2.14 (dqd, *J* = 7.6, 7.3, 1.2 Hz, 2H), 1.76-1.64 (m, 4H), 1.59-1.51 (m, 3H), 1.35 (quint, *J* = 7.6 Hz, 2H), 1.00-0.93 (m, 9H); ^13^C-NMR (100 MHz, CDCl_3_) δ_C_: 210.0, 176.2, 174.5, 173.4, 139.3, 132.3, 125.2, 50.9, 41.1, 36.2, 34.2, 31.1, 29.2, 29.1, 27.1, 25.3, 24.9, 22.9, 21.8, 21.2, 20.6, 14.2; IR (film) cm^−1^: 3314, 2870, 2958, 2931,1685, 1631, 1540, 1439, 1366, 1231, 1057, 670; HRMS (ESI, negative) *m/z* [M–H]^−^ Calcd. for C_22_H_34_NO_4_^−^: 376.2493, found: 376.2478.

### Synthesis of dn-*iso*-OPDA-Ile

To a solution of dn-*iso*-OPDA (51.6 mg, 0.20 mmol) and Et_3_N (85.0 µL, 0.61 mmol) in THF (2.0 mL) was added ethyl chloroformate (40.0 µL, 0.42 mmol) at 0 °C under an argon atmosphere. After stirring at room temperature for 2 h, the above mixture was transferred to a solution of L-isoleucine (123 mg, 0.93 mmol) and diisopropylethylamine (200 µL, 1.17 mmol) in H_2_O (2.0 mL) and the resulting mixture was stirred for 3 h at room temperature. The reaction mixture was then acidified with 1 M aq. HCl and extracted with CHCl_3_. The combined organic layer was washed with brine, dried over Na_2_SO_4_, and concentrated under reduced pressure. The residue was purified by silica gel medium-pressure chromatography (CHCl_3_/MeOH/AcOH = 99/1/0.1 to CHCl_3_/MeOH/AcOH = 90/10/0.1) to give dn-*iso*-OPDA-Ile (66.5 mg, 89%) as a pale yellow oil. The obtained dn-*iso*-OPDA-Ile (17.3 mg) (Supplemental Figure S1 and S3) was further purified by RP-HPLC (ODS-HG-5 column: F 20 × 250 mm, flow rate 8.0 mL/min, detection 220 nm, eluent: MeOH/H_2_O/AcOH = 80/20/0.1, t_R_ = 11.4 min) to give dn-*iso*-OPDA-Ile (11.8 mg) as a colorless oil.

[α]_D_^28^ = +19.4 (c 0.52, CHCl_3_). ^1^H-NMR (400 MHz, CDCl_3_) δ_H_: 6.00 (dd, *J* = 8.2, 0.9 Hz, 1H), 5.36 (dtt, *J* = 10.5, 7.2, 1.3 Hz, 1H), 5.19 (dtt, *J* = 10.5, 7.2, 1.4 Hz, 1H), 4.60 (dd, *J* = 8.2, 4.9 Hz, 1H), 2.92 (d, *J* = 7.2 Hz, 2H), 2.49-2.48 (m, 2H), 2.43 (t, *J* = 7.5 Hz, 2H), 2.38-2.36 (m, 2H), 2.26 (t, *J* = 7.5 Hz, 2H), 2.14 (dq, *J* = 7.6, 7.2 Hz, 2H), 1.98-1.92 (m, 1H), 1.68 (quint, *J* = 7.5 Hz, 2H), 1.55 (quint, *J* = 7.5 Hz, 2H), 1.53-1.45 (m, 1H), 1.36 (quint, *J* = 7.5 Hz, 2H), 1.25-1.17 (m, 1H), 0.98 (t, *J* = 7.3 Hz, 3H), 0.98-0.92 (m, 6H); ^13^C-NMR (100 MHz, CDCl_3_) δ_C_: 209.9, 174.9, 174.5, 173.1, 139.3, 132.3, 125.2, 56.4, 37.5, 36.3, 34.2, 31.1, 29.7, 29.2, 27.1, 25.4, 25.1, 21.2, 20.6, 15.5, 14.2, 11.6; IR (film) cm^-1^: 3308, 2931, 1722, 1629, 1542, 1459, 1358, 993; HRMS (ESI negative) *m/z* [M–H]^−^ Calcd. for C_22_H_34_NO_4_^−^: 376.2493, found: 376.2461.

### Synthesis of dn-*iso*-OPDA-Val

To a solution of dn-*iso*-OPDA (50.5 mg, 0.19 mmol) and Et_3_N (80.0 µL, 0.58 mmol) in THF (2.0 mL) was added ethyl chloroformate (40.0 µL, 0.42 mmol) at 0 °C under an argon atmosphere. After stirring at room temperature for 2 h, the above mixture was transferred to a solution of L-valine (111 mg, 0.95 mmol) and diisopropylethylamine (194 µL, 1.17 mmol) in H_2_O (2.0 mL) and the resulting mixture was stirred for 3 h at room temperature. The reaction mixture was then acidified with 1 M aq. HCl and extracted with CHCl_3_. The combined organic layer was washed with brine, dried over Na_2_SO_4_, and concentrated under reduced pressure. The residue was purified by silica gel medium-pressure chromatography (CHCl_3_/MeOH/AcOH = 99/1/0.1 to CHCl_3_/MeOH/AcOH = 90/10/0.1) to give dn-*iso*-OPDA-Val (55.9 mg, 81%) as a pale yellow oil. The obtained dn-*iso*-OPDA-Val (11.5 mg) (Supplemental Figure S1 and S4) was further purified by RP-HPLC (ODS-HG-5 column: F 20 × 250 mm, Flow rate 8.0 mL/min, detection 220 nm, eluent: MeOH/H_2_O/AcOH = 75/25/0.1, t_R_ = 11.3 min) to give dn-*iso*-OPDA-Val (8.0 mg) as a colorless oil.

[α]_D_^26^ = +17.5 (c 0.86, CHCl_3_). ^1^H-NMR (400 MHz, CDCl_3_) δ_H_: 5.95 (d, *J* = 8.3 Hz, 1H), 5.36 (dtt, *J* = 10.5, 7.3, 1.8 Hz, 1H), 5.20 (dtt, *J* = 10.5, 7.3, 1.4 Hz, 1H), 4.57 (dd, *J* = 8.3, 5.0 Hz, 1H), 2.92 (d, *J* = 7.1 Hz, 2H), 2.50-2.48 (m, 2H), 2.44 (t, *J* = 7.6 Hz, 2H), 2.38-2.36 (m, 2H), 2.28 (t, *J* = 7.6 Hz, 2H), 2.25-2.20 (m, 1H), 2.14 (dq, *J* = 7.3, 7.4 Hz, 2H), 1.69 (quint, *J* = 7.6 Hz, 2H), 1.56 (quint, *J* = 7.6 Hz, 2H), 1.36 (quint, *J* = 7.6 Hz, 2H), 1.00-0.94 (m, 9H); ^13^C-NMR (100 MHz, CDCl_3_) δ_C_: 210.0, 174.9, 174.6, 173.4, 139.3, 132.3, 125.2, 57.0, 36.4, 34.2, 31.1, 30.9, 29.7, 29.2 (2C), 27.1, 25.4, 21.2, 20.6, 19.0, 17.7, 14.2; IR (film) cm^−1^: 3328, 3011, 2963, 2933, 2871, 1729, 1684, 1632, 1541, 1464, 1360, 1205; HRMS (ESI, negative) *m/z* [M-H]^−^ Calcd. for C_21_H_32_NO_4_^−^: 362.2337, found: 362.2336.

### Analysis of dn-*iso*-OPDA and dn-*iso*-OPDA conjugated with His, Glu, Gln, Val, Leu and Ile

Endogenous dn-*iso*-OPDA and dn-iso-OPDA conjugated with His, Glu, Gln, Val, Leu and Ile were analyzed using high performance liquid chromatography-electrospray-high-resolution accurate mass spectrometry (HPLC-ESI-HRMS).

As previously described (Chini et al. 2021; Chini et al. 2023), these molecules were extracted from approximately 0.1g of vegetal material, ground to a powder in a mortar with liquid N_2_. 1 mL of a precooled (-20°C) mixture of MeOH/H_2_O/formic acid (90:9:1, v/v/v) containing 2.5mM of sodium diethyldithiocarbamate and 10µL of a stock solution of the deuterium-labeled internal standards in methanol was added to the frozen vegetal material. The extraction was developed by two steps of shaking and centrifugation. First, the extraction was performed shaking the samples during 60 min at 2000 rpm at room temperature using a Multi Reax shaker (Heidolph Instruments, Schwabach, Germany). Solids were separated by centrifugation at 13.000 rpm for 10 min using a Biofuge pico Centrifuge (Thermo Fisher Scientific, Waltham, Massachusetts, USA). The extraction was repeated adding 0.5mL to the initial solid and shaking during 20 min. 1mL of the combined supernatants from the two extraction procedures was evaporated (40°C) until all the solvent disappeared (RapidVap Evaporator (Labconco Co., Kansas City, MO). The residue was re-dissolved with 0.25 mL of a mixture of methanol/acetic acid (0.133%) (40:60, v/v) and centrifuged at 20000 RCF using a Sigma 4-16K Centrifuge (Sigma Laborzentrifugen gmbH, Osterode am Harz, Germany) before the transfer to an injection vial.

Chromatographic separation of the molecules was performed by a Dionex Ultimate 3000 UHPLC System. A reverse-phase column (Synergi 4 mm Hydro-RP 80A, 150 x 2 mm; Phenomenex, Torrance, CA) was applied. A linear gradient of methanol (A), water (B) and 2% acetic acid in water (C) was used: 38% A for 3 min, 38% to 96% A in 12 min, 96% A for 2.5 min and 96% to 38% A in 0.5 min, followed by a stabilization time of 4 min. The percentage of C remains constant at 4%. The flow rate was 0.30 mL min^-1^, the injection volume was 20 µL and column and sample temperatures were 35 and 15 °C.

The detection and quantification were performed using an Exploris 120 mass spectrometer (ThermoFisher Scientific) equipped with an OptaMax NG ion source, a C-Trap, an Ion-Routing Multipole, a High Field Orbitrap Mass Analyzer with a quadrupole mass filter. The instrumental analysis parameters are reported in Supplementary Table S2. The procedure was developed using a Product Ion Scan Experiment in the negative-ion mode for dn-iso-OPDA and a Full Scan experiment in the negative-ion mode for the conjugates, employing multilevel calibration curves with the internal standard (D-dn-OPDA) for dn-iso-OPDA and with external standards for the conjugates. Resolution was set at 60.000 FWHM in both experiments, Q1 Resolution (m/z) at 1.2 in the Product Ion experiment, AGC target Standard, Maximum injection time mode Auto and RF Lens at 70%. Three fragments were analyzed for dn-iso-OPDA: the fragment ion with the highest intensity (fragment 1) was used for quantification, and the other two (fragment 2 and fragment 3) were used for the confirmation of the molecular identity. For D-dn-OPDA, only the fragment ion of the highest intensity was analyzed. The Absolute Collision Energy (CE) for dn-iso-OPDA and D-dn-OPDA is 13 V. For the conjugates only the parent ion was analyzed (Supplementary Table S3). A mass tolerance of 5 ppm was accepted. TraceFinder 5.3 EFS software was used for data processing and instrument control.

### Snail assays

Petri dishes (9 cm in diameter) were prepared by drawing a straight line across the back to divide them into two equal halves. Ten Tak-1 gemmae were planted on one half, while ten gemmae of the comparative genotypes (Mp*coi1-2*, Mp*ilr1-1^ge^* or Mp*ilr1-2^ge^*) were planted on the other. On day 10, plants were photographed prior to the challenge, and then infested with eight snails *Helix aspersa* (approximately 1 cm in size) per plate (Espinosa et al., 2026). After 24 hours of feeding, the snails were removed and the plates were photographed again. Region identification was performed with AI-assisted image analysis (color-threshold segmentation and image registration, Claude, Anthropic), with manual verification against the original photographs. ImageJ software was utilized to measure the thallus surface area before and after snail infestation, from which the remaining thallus area percentage was calculated.

### Insect assays

For the insect assays, ten plants of each genotype were cultivated on plates for 4 weeks before introducing 12 first-instar *S. exigua* larvae (Entocare, Wageningen, The Netherlands) per plate, with 5 to 10 replicate plates utilized per genotype. Following a 7- to 8-day feeding period, the larvae were individually collected and weighed using a precision balance. This experiment was independently replicated four times with consistent results. Data are presented as the mean ± SD.

### Metabolite analysis after snail and insect infection

Following infestation, thalli were collected, immediately submerged in liquid nitrogen, and ground into a fine powder. The pulverized tissue was mixed with 1 mL of 90% methanol, and a 900 µl aliquot of the supernatant was collected. The liquid was then completely evaporated using a vacuum concentrator (SpeedVac) and stored at −20 °C until metabolite analysis.

### Accession numbers

Sequence data from this article can be found in the GenBank/EMBL data libraries under accession numbers: MpCOI1/Mp2g26590, MpILR1/Mp1g20090 and MpGH3A/Mp6g07600

## Results

### MpILR1 is required for accumulation of specific dn-*iso*-OPDA conjugates

To investigate whether dn-*iso*-OPDA amino acid conjugates undergo hydrolysis and thereby re-enter the active hormone pool in *Marchantia polymorpha*, we generated CRISPR-Cas9 loss-of-function mutants of Mp*ILR1*, the only predicted ILR1/ILL-family amidohydrolase encoded in the *M. polymorpha* genome. We isolated two independent alleles, Mp*ilr1-1^ge^* and Mp*ilr1-2^ge^*, carrying deletions of 932 and 952 nucleotides, respectively (Supplementary Fig. S5). Both deletions removed most of the first exon and introduce a premature stop codon in both cases (Supplementary Fig. S1). Next, we analyzed accumulation of dn-*iso*-OPDA amino acid conjugates in wild-type (WT) and Mp*ilr1* plants using ultra-performance liquid chromatography coupled to tandem mass spectrometry (UPLC-MS/MS). Mechanical wounding induced the major conjugates dn-*iso*-OPDA-His, -Glu and -Gln to similar levels in WT and both Mp*ilr1* alleles (Fig. 1A). In contrast, less abundant conjugates with the predicted masses of dn-*iso*-OPDA-Val and dn-iso-OPDA-Leu/Ile accumulated to significantly higher levels in both Mp*ilr1* alleles than in WT plants after wounding (Fig. 1B). Because dn-*iso*-OPDA-Leu and -Ile have identical masses and could not be resolved under our UPLC–MS/MS conditions, their signals were jointly quantified. To confirm conjugate identities, we synthesized authentic dn-*iso*-OPDA-Val, -Leu and -Ile standards (Supplementary Fig. S1 and S2).

**Figure. 1.**
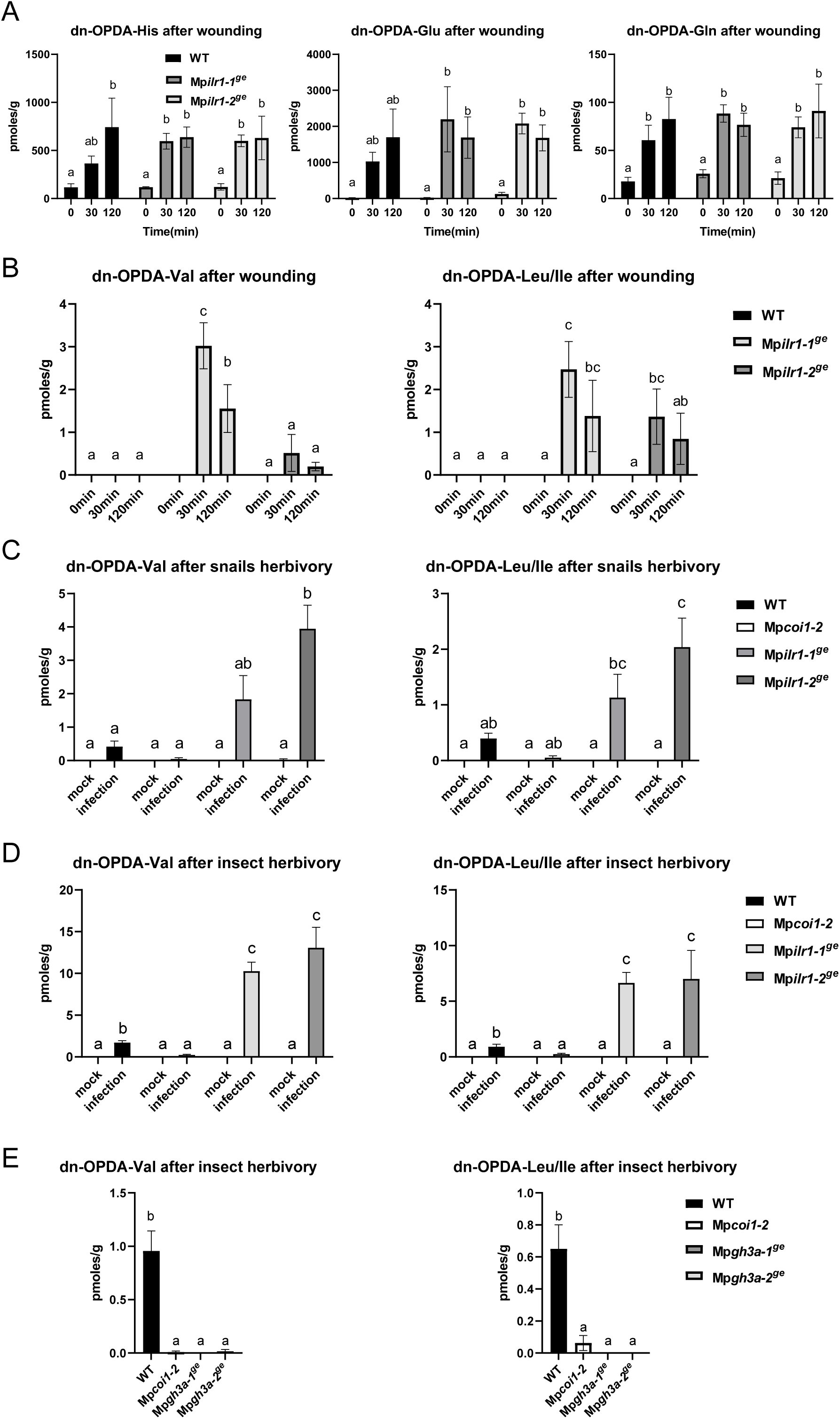
Accumulation of dn-*iso*-OPDA conjugates upon stress in *M. polymorpha*. Time-course accumulation of dn-*iso*-OPDA conjugates [pmol/fresh weight (g)] in WT (Tak-1) *Marchantia* plants, Mp*coi1-2*, Mp*ilr1-1^ge^* or Mp*ilr1-2^ge^* mutants after wounding (**A** and **B**), or gastropod (**C**) and insect (**D**) herbivory challenge. Data shown as mean ± SD of three biological replicates. Experiments were repeated three times with similar results. **E**) Accumulation of dn-*iso*-OPDA-Val and -Leu/Ile after insect herbivory infection in WT, Mp*coi1-2* and Mp*gh3a^ge^* mutants. Letters indicate significant different samples according to the one-way ANOVA/Tukey HSD post hoc test (P < 0.05).

Quantification using authentic standards showed that the selective accumulation of hydrophobic conjugates in *Mpilr1* mutants was not restricted to mechanical damage. After feeding by either *Helix aspersa* or *Spodoptera exigua*, dn-*iso*-OPDA-Val and -Leu/Ile were consistently more abundant in *Mpilr1* than in WT plants, whereas the levels of the His, Glu and Gln conjugates did not differ between genotypes (Fig. 1C-D; Supplementary Fig. S6). These results indicate that MpILR1 is required for hydrolysis of a defined subset of dn-*iso*-OPDA conjugates. However, direct enzymatic assays will be required to confirm substrate specificity and kinetic parameters. In addition, Mp*gh3a-1^ge^* plants failed to accumulate dn-*iso*-OPDA-Val and -Leu/Ile following *S. exigua* herbivory (Fig. 1E), indicating that their production requires MpGH3A. Thus, MpGH3A generates several dn-*iso*-OPDA amino acid conjugates, whereas MpILR1 preferentially promotes turnover of the Val and Leu/Ile derivatives.

### MpILR1-mediated hydrolysis of specific dn-*iso*-OPDA conjugates modulates dn-OPDA responses in *M. polymorpha*

The enhanced accumulation of dn-*iso*-OPDA-Val and -Leu/Ile in Mp*ilr1* plants suggested that these compounds might function as hydrolyzable precursors of the bioactive hormone, despite their low abundance. Therefore, we tested the response of WT, Mp*ilr1*, and Mp*coi1-2* plants to conjugate exogenous treatments. dn-*iso*-OPDA-Val and -Leu significantly inhibited WT growth, whereas dn-*iso*-OPDA-Ile did not produce a substantial growth reduction under the conditions used (Fig. 2A-B). Growth inhibition by both dn-*iso*-OPDA-Val and -Leu was strongly reduced or absent in the OPDA-insensitive Mp*coi1-2* plants, demonstrating that these responses require MpCOI1 (Fig. 2A-B). Loss of Mp*ILR1* function also reduced sensitivity to dn-*iso*-OPDA-Val and -Leu (Fig. 2A-B). In contrast, treatment with dn-*iso*-OPDA-His, -Glu or -Gln did not significantly inhibit growth in either WT or Mp*ilr1-1^ge^*plants (Fig. 3A-B).

**Figure 2:**
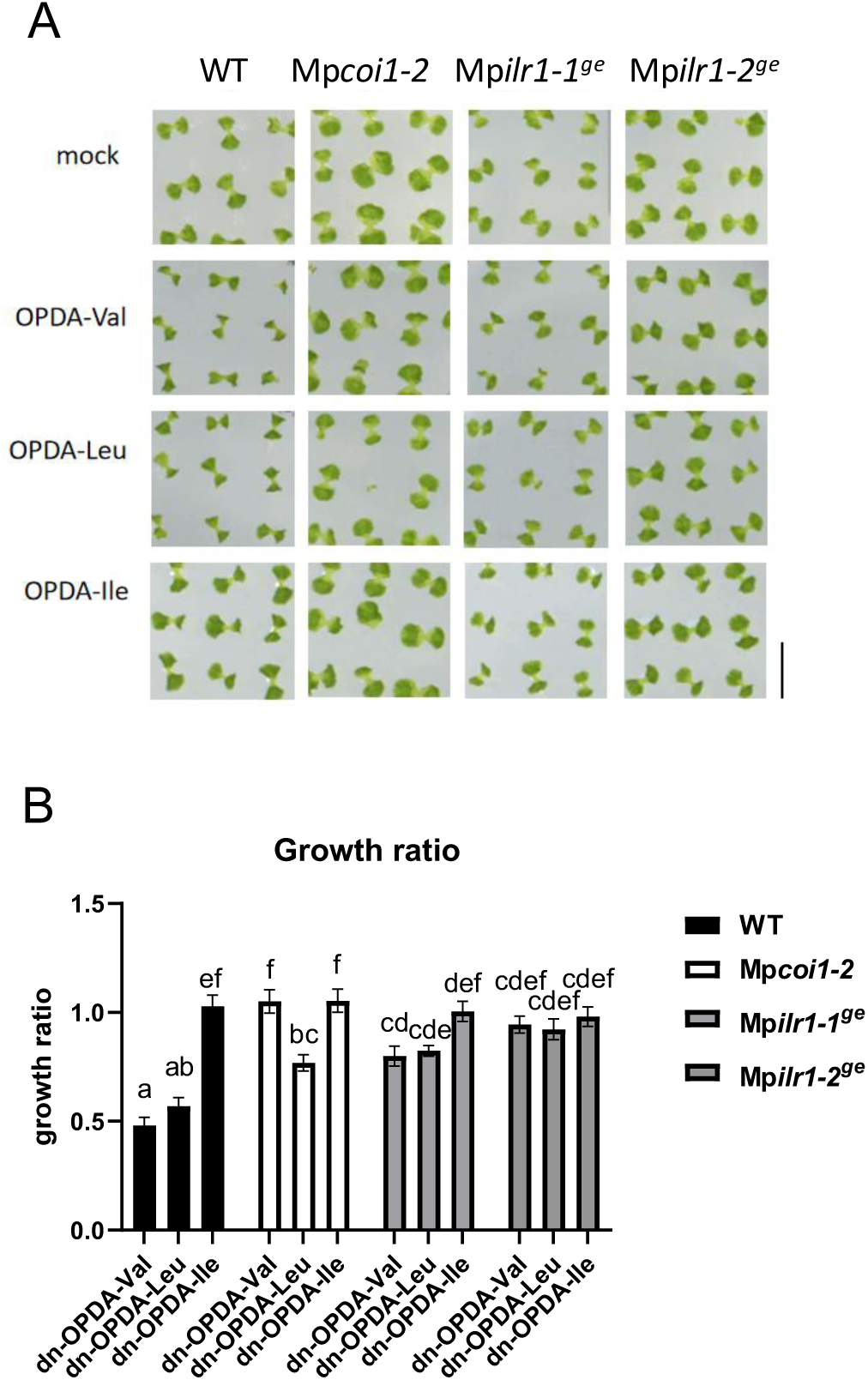
Effect of OPDA-VAL, OPDA-LEU and OPDA-ILE on the growth of WT and Mp*ilr1* mutant plants. **A**) Images of the plants grown for 13 days in mock plates or in the presence of the OPDA-amino acids. **B**) Quantification of the growth inhibition induced by OPDA-amino acids in wild-type Tak-1 (WT), Mp*coi1-2* and Mp*ilr1* mutant plants grown for 13 days in absence (mock) and presence of 10μM OPDA-amino acids. Data are shown as mean ± SD of three biological replicates, 12 plants per each replicate. Experiments were repeated three times with similar results. Letters indicate significant different samples according to the one-way ANOVA/Tukey HSD post hoc test (P < 0.05).

**Figure 3.**
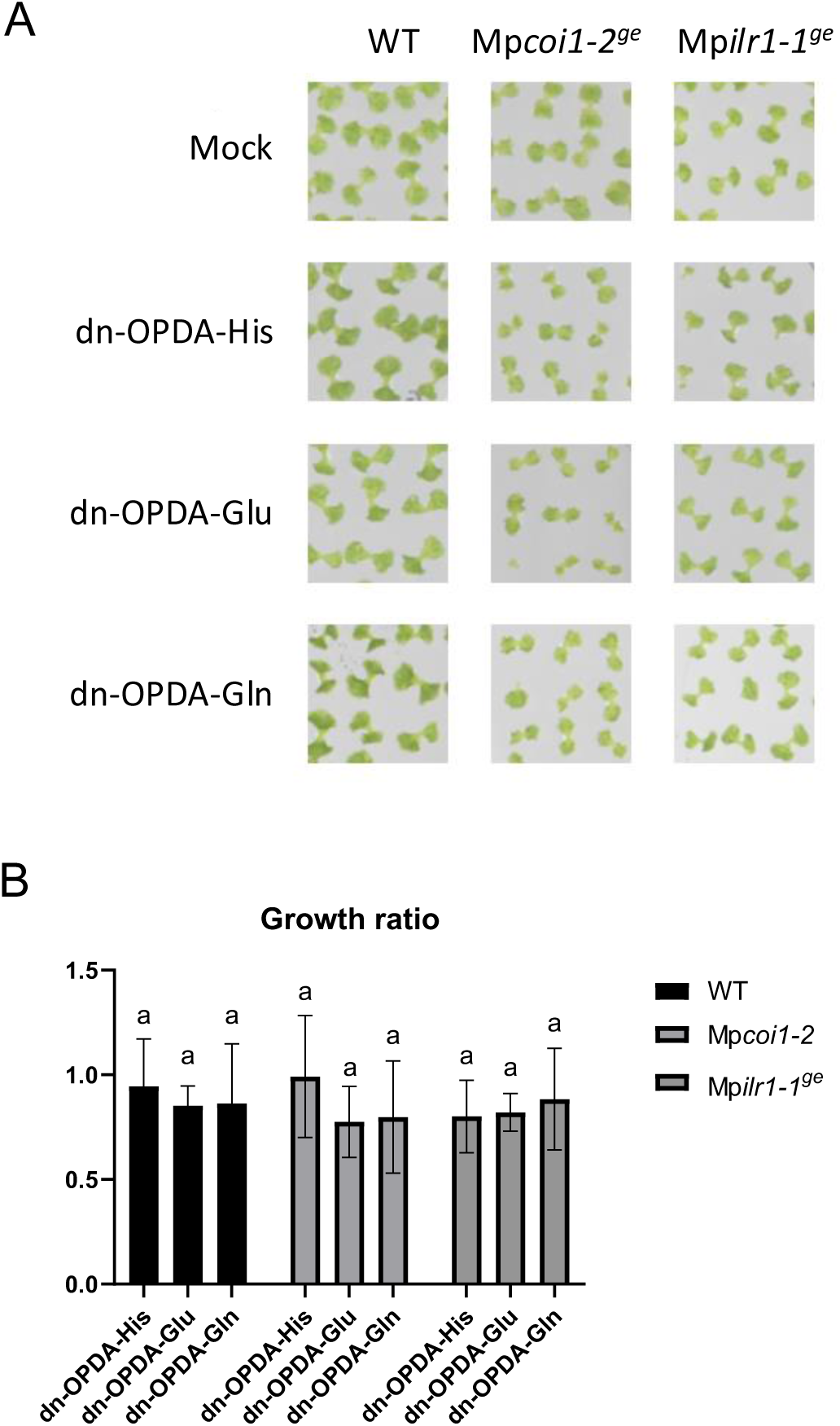
OPDA-His, OPDA-Glu and OPDA-Gln do not inhibit growth in WT and Mp*ilr1* mutant plants. **A**) Images of the plants grown for 13 days in mock plates or in the presence of the OPDA-amino acids. **B**) Quantification of the growth inhibition induced by OPDA-amino acids in wild-type Tak-1 (WT), Mp*coi1-2* and Mp*ilr1* mutant plants grown for 13 days in absence (mock) and presence of 10μM OPDA-amino acids. Data are shown as mean ± SD of three biological replicates, 12 plants per each replicate. Letters indicate significant different samples according to the one-way ANOVA/Tukey HSD post hoc test (P<0.05).

We further tested whether dn-*iso*-OPDA conjugates affect transcriptional activation of the dn-OPDA pathway. In WT plants, treatment with dn-*iso*-OPDA-Val, -Leu or -Ile induced the expression of the dn-OPDA marker Mp*PAT* and Mp*DIR* relative to mock control (Fig. 4A). In contrast, dn-*iso*-OPDA-His, -Glu or -Gln did not generally induce the marker gene expression. To determine whether the transcriptional activation induced by dn-*iso*-OPDA conjugates requires MpILR1, we treated Mp*ilr1* mutants with the hydrolyzable conjugate subset, -Val, -Leu or -Ile. Induction of dn-*iso*-OPDA marker genes was absent in Mp*coi1-2* plants and significantly weaker in Mp*ilr1* mutants than in WT plants (Fig. 4B). Thus, MpILR1-dependent turnover of specific dn-*iso*-OPDA amino acid conjugates contributes to activate MpCOI1-dependent growth and gene-expression responses.

**Figure 4.**
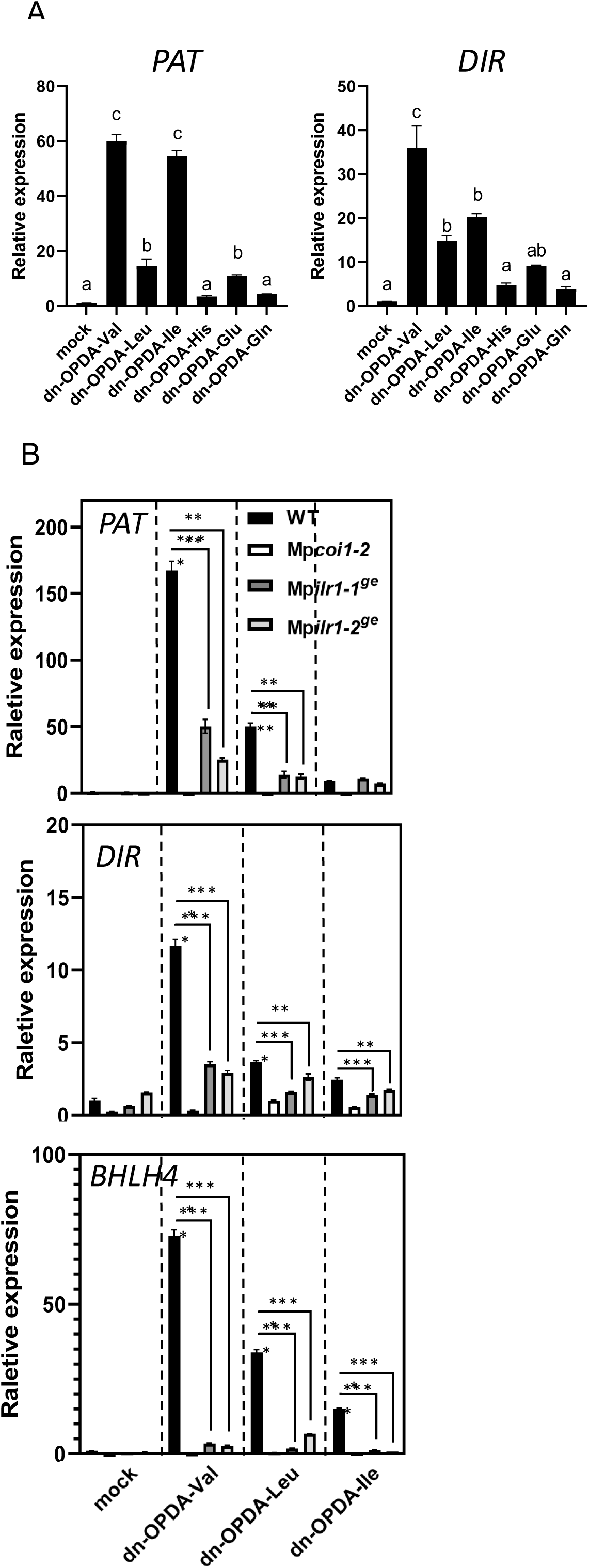
Transcriptional analyses of jasmonate markers. **A**) Expression level of bioactive marker gene *PATATIN* and *DIRIGIN* in WT Tak-1 plats after different dn-OPDA-amino acid treatments. Letters indicate significant different samples according to the one-way ANOVA/Tukey HSD post hoc test (P<0.05). **B**) Expression level of bioactive marker gene *PATATIN*, *DIRIGIN* and *BHLH4* after 2 hours of dn-OPDA-Val, dn-ODPA-Leu and dn-OPDA-Ile treatment in WT, Mp*coi1-2*, and Mp*ilr1* plants. Statistical significance was determined using unpaired two-tailed Student’s t-test. Asterisks denote significance levels: *P < 0.05, **P < 0.01, ***P < 0.001, ****P < 0.0001; ns, not significant.

### MpILR1-dependent jasmonate homeostasis regulates defense against gastropod herbivory

Next, we investigated whether MpILR1-mediated conjugate turnover contributes to regulate jasmonate-mediated defenses. Because dn-*iso*-OPDA/MpCOI1 signaling pathway contributes to defense against gastropod feeding in *M. polymorpha*, we examined whether loss of MpILR1 activity affects snail resistance (Schweizer et al., 2025). In two-choice feeding assays, snails displayed a significant feeding preference for Mp*coi1-2* relative to WT, confirming the requirement for the dn-*iso*-OPDA/MpCOI1 pathway in gastropod defense (Fig. 5A-B) (Schweizer et al., 2025; Espinosa-Cores et al., 2026). Snails also showed a significant feeding preference for Mp*ilr1* mutants over WT plants grown on the same plate, indicating that MpILR1 contributes to full resistance to gastropod herbivory (Fig. 5A-B).

**Figure 5:**
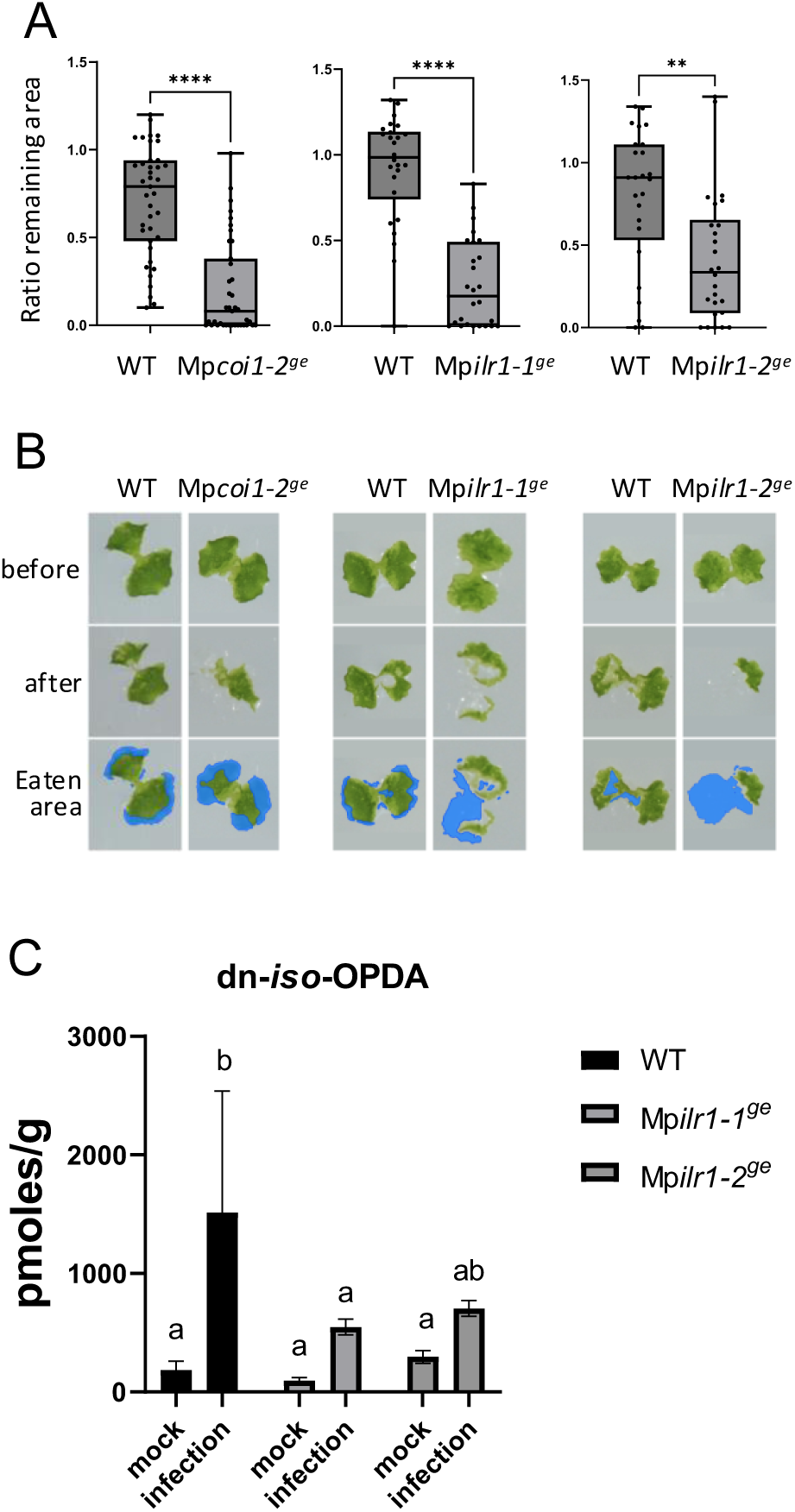
Mp*ILR1* regulates defense against snail herbivory. Snail infection was designed as a two-choice experiment and carried out on 2 week-old plants, comparing WT Tak-1 with one individual mutant line, Mp*coi1-2*, Mp*ilr1-1^ge^* or Mp*ilr1-2^ge^* mutants. Two-choice assays were performed at the same time. Quantification of thallus area change ratio was defined by area quantification after and before the snail feeding assay (n=39 to 52) (**A**). Asterisks indicate significant differences compared to the untreated control WT Tak-1 plants evaluated by tests (* P<0.1). Experiments were repeated two times with similar results. **B**) Plant pictures before and 24 hours after snail infection. The blue area represents the snail’s eating area. **C**) Accumulation of dn-*iso*-OPDA conjugates after snail infection in WT and Mp*ilr1* plants. Letters indicate significant different samples according to the one-way ANOVA/Tukey HSD post hoc test (P < 0.05).

To investigate whether this diminished defense was associated with altered hormone accumulation, we measured dn-*iso*-OPDA levels following snail feeding. Gastropod herbivory significantly increased dn-*iso*-OPDA accumulation in WT plants, whereas this induction was partially reduced in Mp*ilr1* mutant alleles (Fig. 5C). Notably, although dn-*iso*-OPDA-Val, -Leu and -Ile constituted a minor fraction of the total dn-OPDA pools, they show a substantial effect to dn-*iso*-OPDA homeostasis and gastropod resistance. How this important effect is mechanistically achieved is an important question that will require further investigation.

Together with the accumulation of dn-*iso*-OPDA-Val and -Leu/Ile in Mp*ilr1* plants, these results support the conclusion that MpILR1-mediated conjugate turnover contributes to maintaining bioactive dn-*iso*-OPDA levels and to the full activation of MpCOI1-dependent anti-herbivore defense in *M. polymorpha*. They further suggest that a reversible cycle of dn-*iso*-OPDA conjugation and hydrolysis evolved as a mechanism for hormonal regulation during plant colonization of land.

## Discussion

In Arabidopsis, hormone homeostasis depends on a bidirectional cycle of conjugation, mediated by GH3-type acyl-acid amido synthetases, and deconjugation or hydrolysis, mediated by ILR/ILL-type amidohydrolases (Zhang et al., 2016; Jez, 2022). For example, OPDA- and JA-amino acid conjugates are rapidly generated upon stress and either serve as transient storage forms (OPDA-aa) or as bioactive COI1-JAZ ligands (JA-Ile and related JA-aa conjugates) that are themselves subject to rapid turnover, thereby buffering cis-OPDA and JA-Ile pools (Yan et al., 2016; Marquis et al., 2020; Široká et al., 2025). In bryophytes, only the conjugating branch of this cycle had previously been genetically characterized: MpGH3A-dependent conjugation of dn-*iso*-OPDA to amino acids diminishes the bioactive dn-*iso*-OPDA pool in *M. polymorpha* (Liang et al., 2025). These findings indicated that conjugation-based inactivation regulates attenuation of dn-*iso*-OPDA signaling; however, whether such conjugates represent terminal catabolic sinks or components of a reversible cycle, analogous to the angiosperm ILR/ILL pathway, remained unresolved.

Here, we show that a previously uncharacterized and low abundant subset of dn-*iso*-OPDA-amino acid conjugates in *M. polymorpha* participates in a reversible conjugation-hydrolysis cycle that dynamically buffers the bioactive dn-*iso*-OPDA pool. To our knowledge, this is the first evidence that dn-*iso*-OPDA-amino acid conjugates function as a dynamic hormone reservoir in a bryophyte species, rather than serving exclusively as irreversible catabolic end products.

Our data indicate that only a defined subset of dn-*iso*-OPDA-amino acid conjugates is efficiently hydrolyzed, supporting a model in which conjugate nature dictates distinct metabolic fates ranging from metabolically inert (hydrophilic and polar Gln, Glu or His) end products to readily mobilizable hormone reservoirs (hydrophobic Val, Ile and Lue). Within this homeostatic circuit, impaired conjugation, as previously described for Mp*gh3a* mutants, elevates bioactive dn-*iso*-OPDA levels and thereby enhances COI1-dependent signaling and downstream defense responses (Liang et al., 2025). MpILR1, conversely, functions as an amidohydrolase mediating the deconjugation branch of this cycle, selectively cleaving a defined subset of dn-*iso*-OPDA-amino acid conjugates to restore free, bioactive dn-*iso*-OPDA from an inactive reservoir. Consistent with this role, loss-of-function Mp*ilr1* mutants accumulate specific dn-*iso*-OPDA-amino acid conjugates and show a corresponding reduction in stress-induced release of bioactive dn-*iso*-OPDA. While MpILR1 is the only annotated ILR1/ILL ortholog, the low abundance accumulation of specific dn-*iso*-OPDA-amino acid conjugates in Mp*ilr1* mutants raise the possibility that the timing used in our assays may have missed the stronger accumulation peaks. Additionally, functional redundancy with other amidohydrolase enzymes contributing to conjugate turnover cannot be discarded. At the transcriptional and physiological levels, impaired MpILR1 activity results in significantly attenuated induction of dn-*iso*-OPDA-responsive genes and reduced growth inhibition following exogenous treatment with dn-*iso*-OPDA-amino acid conjugates. Finally, Mp*ilr1* mutants showed diminished resistance to gastropod herbivory, underscoring the role of MpILR1-mediated hydrolysis in full defense activation.

Notably, MpILR1 hydrolytic activity is selective toward a minor subset of dn-OPDA-amino acid conjugates rather than the most abundant conjugate species generated upon stress. The major conjugate class comprises dn-*iso*-OPDA conjugated to hydrophilic, polar or charged amino acids (His, Glu, Asp or Gln), which appear to function as irreversible catabolic end products and thus constitute a stable sink for the active hormone. By contrast, the minor conjugate class, formed with hydrophobic, non-polar amino acids (Val, Ile or Leu), can be efficiently hydrolyzed back to free hormone. This distinction between these conjugate pools suggests that MpILR1-mediated turnover of aliphatic dn-*iso*-OPDA conjugates provides a mechanism for fine-tuning hormone availability without interfering with the inactivation route governed by the more abundant, irreversibly conjugated species.

Overall, the MpILR1-mediated regulation of dn-*iso*-OPDA homeostasis described here provides a previously missing layer of jasmonate regulation in bryophytes and reveals that even low-abundance conjugate pools can exert a substantial control over the accumulation and signaling output of bioactive dn-*iso*-OPDA. This apparent discrepancy between conjugates pool size and regulatory impact can be reconciled through several non-exclusive mechanistic considerations. For example, steady-state metabolite abundance does not necessarily reflect flux capacity: even small dn-*iso*-OPDA-aa pools may turn over rapidly through cycles of conjugation and deconjugation, such that cumulative flux over a stress episode substantially exceeds instantaneous pool size. This principle is well established in Arabidopsis, where OPDA-aa conjugates accumulate to only a few pmol/g after wounding, yet isotope-labeling experiments demonstrate that OPDA-Ile and OPDA-Val are rapidly hydrolyzed and converted into JA and JA-Ile, confirming their function as high-flux intermediates that substantially contribute to jasmonate production (Široká et al., 2025). By analogy, MpGH3A and MpILR1 likely support a high-flux dynamic cycle in which dn-*iso*-OPDA molecules dynamically enter and exit the conjugated state, enabling a relatively small conjugate pool to exert influence exceeding expected levels.

Moreover, the regulatory capacity of dn-*iso*-OPDA conjugates likely depends on local and temporal accumulation rather than bulk tissue concentration. dn-*iso*-OPDA and its conjugates are unlikely to be uniformly distributed; instead, their synthesis and hydrolysis may instead occur within specific subcellular compartments and cell types. Single-cell RNA sequencing of *M. polymorpha* shows that Mp*GH3A* and Mp*ILR1* are both highly expressed in air-pore and oil-body cells (Wang et al., 2023), suggesting that dn-*iso*-OPDA conjugates may preferentially accumulate within these specific cell types. In addition, Mp*GH3A* and Mp*ILR1* were specifically expressed in distinct clusters identified by single-nucleus RNA sequencing of Marchantia reproductive structures, antheridiophores and archegoniophores (Zeng et al., 2026). Furthermore, angiosperm GH3 and ILR/ILL enzymes acting on auxin and auxin-conjugates localize in the cytosol, endoplasmic reticulum and nucleus, positioning them to control hormone availability close to or precisely where receptor complexes and transcriptional regulators reside (Sanchez Carranza et al., 2016; Helusová et al., 2025; Liu et al., 2025; Včelařová et al., 2026). MpILR1 may be subject to comparable spatial regulation, such that localized deconjugation near the MpCOI1-MpJAZ co-receptor, or within microdomains of elevated dn-*iso*-OPDA concentration, could impose regulatory control that is not evident from whole-tissue metabolomic analyses. Recent advances in our understanding of auxin subcellular compartmentalization, and its close integration with auxin conjugation and hydrolysis, provide a framework for experimentally studying the subcellular distribution of metabolites, including dn-OPDA-derivates (Včelařová et al., 2026).

Our finding that MpILR1 acts on a dn-*iso*-OPDA-aa pool to regulate dn-*iso*-OPDA-mediated responses suggests that the conjugation-deconjugation homeostatic mechanism may be an ancient regulatory module that evolved in the common ancestor of all land plants. Although MpILR1 phylogenetic analyses supports its position as the ancestor of tracheophyte hormonal conjugate hydrolases (Campanella et al., 2018; Bowman et al., 2021), functional characterization of ILR1 orthologs in mosses, hornworts and representative vascular plants would strengthen this evolutionary inference. Moreover, this regulatory module has been conserved throughout land plant evolution, even as its functional polarity was inverted in the case of JA-Ile. In Marchantia, which accumulate dn-*iso*-OPDA but lack detectable JA-Ile, GH3-mediated conjugation inactivates the bioactive ligand, while ILR1-mediated hydrolysis restores it, positioning conjugation as the ‘off’ switch and hydrolysis as the ‘on’ switch of hormone signaling. A comparable regulatory configuration is conserved in auxin and *cis*-OPDA homeostasis in angiosperms. This regulation polarity contrasts with that of JA-Ile in euphyllophytes (vascular plant clade encompassing ferns, horsetails and angiosperms), which accumulate JA-Ile as the canonical COI1-JAZ ligand, but not dn-*iso*-OPDA. In these lineages, GH3-mediated conjugation of JA to isoleucine generates the bioactive hormone JA-Ile, whereas ILR-family hydrolysis of JA-Ile attenuates signaling, making conjugation the ‘on’ switch and hydrolysis the ‘off’ switch (Staswick and Tiryaki, 2004; Widemann et al., 2013; Woldemariam et al., 2012; Zhang et al., 2016). This inversion of GH3/ILR1 regulatory polarity in the case of JA-Ile (Chini et al., 2023) indicates that the evolutionary transition of the COI1-JAZ ligand from dn-*iso*-OPDA to JA-Ile was accompanied by functional repurposing of a pre-existing conjugation-hydrolysis module, rather than by the emergence of an entirely novel mechanism for jasmonate homeostasis.

## Author contributions

Conceptualization: R.S. and A.C.; Performed research: W.L., A.M.Z., T.K., R.T, and H.S.; Data analysis: W.L., A.M.Z., R.S., and A.C.; Writing-original draft preparation: A.C.; Manuscript review and editing: all authors; Experimental supervision: M.U., J.M.G-M., R.S., and A.C.; Funding acquisition: R.S. and A.C. All authors have read and agreed to the published version of the manuscript.

## Declaration of Generative AI and AI-assisted technologies in the writing process

During the preparation of this work, the authors used Perplexity and PaperPal in order to assist with language refinement and the development of alternative wording. The author designed the scientific concept and figure structure, reviewed and edited all AI-assisted outputs, and takes full responsibility for the content, accuracy, and originality of the publication.

## Supplementary Material

**Supplementary Figure S1. Synthesis of dn-iso-OPDA-L-Amino acid conjugates.**

Reagents and conditions: (a) ClCO2Et, Et3N, THF, 0 □; L-amino acid, DIPEA, H2O.

**Supplementary Figure S2. 1H- and 13C-NMR spectra of dn-iso-OPDA-Leu**.

**Supplementary Figure S3. 1H- and 13C-NMR spectra of dn-iso-OPDA-Ile**.

**Supplementary Figure S4. 1H- and 13C-NMR spectra of dn-iso-OPDA-Val**.

**Supplementary Figure S5. Generation of Mp*ilr1^ge^* mutant alleles.**

Scheme of Mp*ILR1* gene of *Marchantia polymorpha* (A); grey blocks represent exons and dark grey block highlights the sequence encoding for the ILR domain. Dashed lines represent deletions in Mp*ILR1^ge^*alleles. Stop codon are highlighted in red. Scheme of MpILR1 protein sequence of WT and Mp*ILR1^ge^* mutant alleles (B); the hyphen symbol “-” indicates a deleted segment.

**Supplementary Figure S6. Accumulation of dn-iso-OPDA-His, -Glu and -Gln after herbivory.**

Accumulation of dn-iso-OPDA conjugates [pmol/fresh weight (g)] in WT (Tak-1) Marchantia plants, Mpilr1-1ge and Mpilr1-2ge mutants after snail (A) and insect herbivory (B). Letters indicate significant different samples according to the one-way ANOVA/Tukey HSD post hoc test (P < 0.05).

**Supplementary Table S1:**
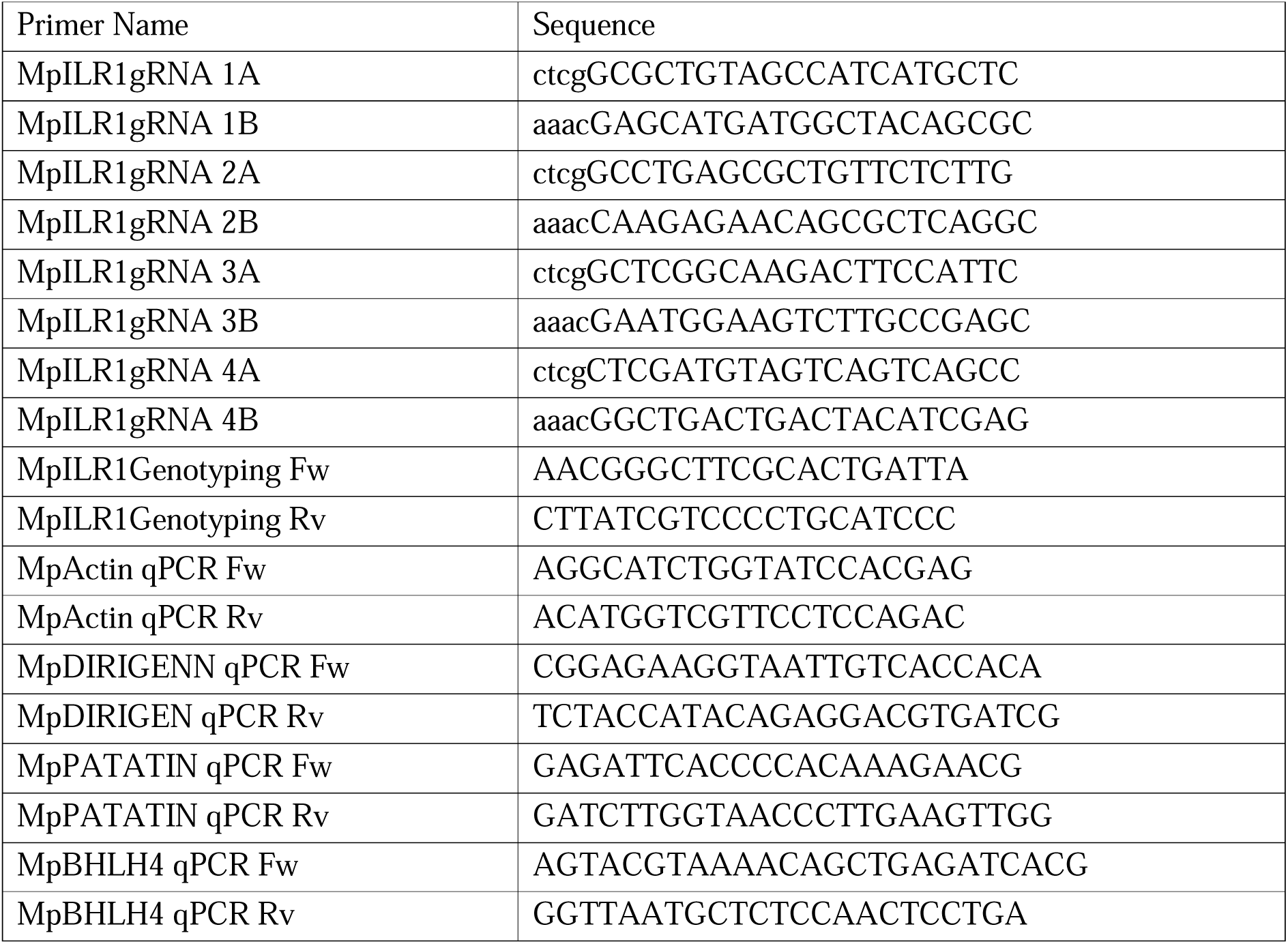
sequences of primers used in this work.

**Supplementary Table S2.**
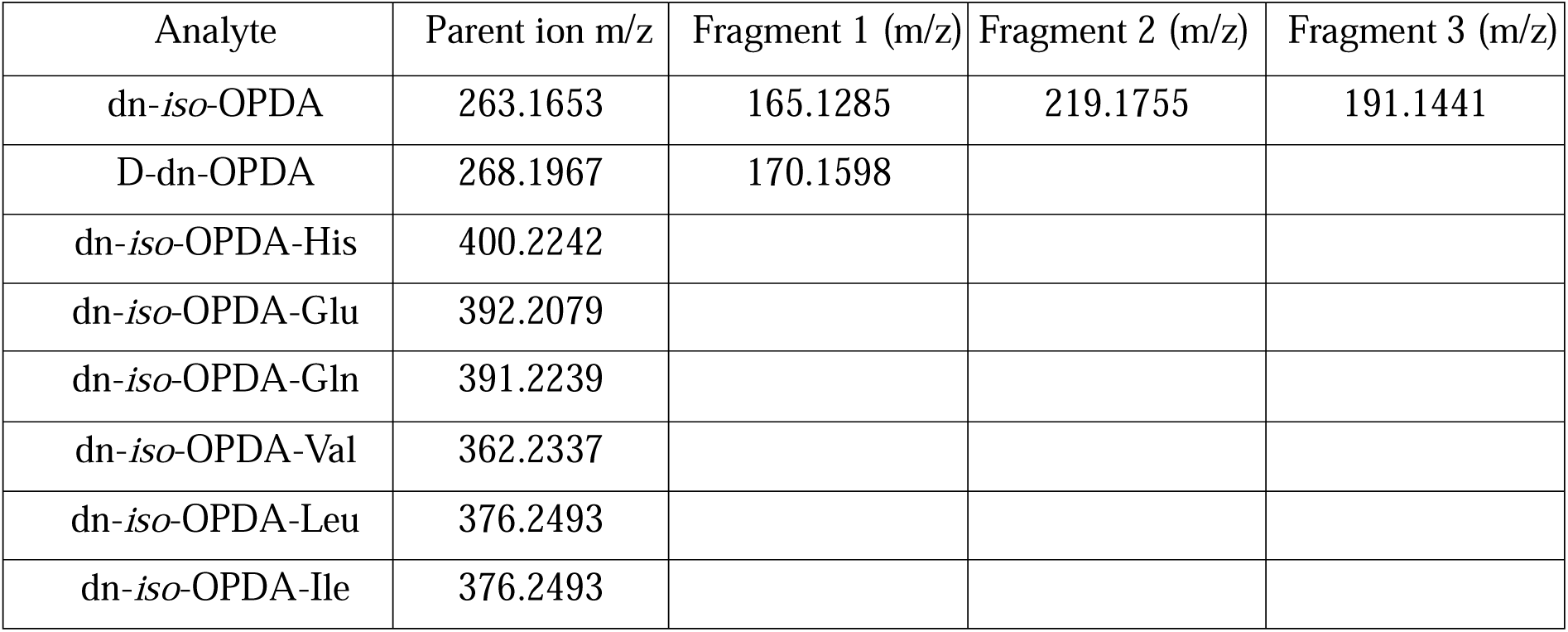
Accurate m/z of the molecules and internal standard, and its principal fragments.

**Supplementary Table S3:** Instrumental parameters used for OptaMax NG ion source.

| Instrumental parameters | Value |
| --- | --- |
| Sheath gas flow rate | 50 au |
| Auxiliary gas flow rate | 10 au |
| Sweep gas flow rate | 1 au |
| Spray voltage | 2900 V |
| Ion transfer tube temperature | 320 °C |
| Vaporizer temperature | 300 °C |

